# Covert attentional tracking with invisible phase tagging

**DOI:** 10.64898/2026.10.21.760697

**Authors:** Kabir Arora, Klara Žinkovič, J. Leon Kenemans, Stefan Van der Stigchel, Surya Gayet, Samson Chota

## Abstract

Attention forms a fundamental component of visual processing. The locus and spread of covert attention reveals how specific locations or features in the environment are preferentially processed, however, tracking this process in a time-resolved fashion is not easy. Rapid Invisible Frequency Tagging (RIFT) is a recent advance in localizing covert attention. This technique uses stimuli that flicker at unique frequencies to invisibly “tag” areas of the visual field. The frequency-specific response elicited offers a measure of visual processing (and covert attention) over time. However, the narrow frequency band compatible with RIFT restricts the number of distinct frequency tags that can be simultaneously used. This limitation can be addressed by using oscillatory phase as an additional tagging feature. Here, we conducted a RIFT-EEG experiment (n=24) with 8 unique tags based on both frequency and phase modulation. We show that modulations in visual processing can be measured at several tagged locations with identical frequency but different phases. This also allowed for the use of combined phase- and frequency-modulated RIFT to quantify the distribution of attention across the visual field. By integrating phase-modulation into invisible frequency tagging studies of attention, we bring the field a step closer towards fine-resolution attentional tracking continuously over time and space, allowing novel insights across various domains of visual cognition.

## Introduction

Attention forms a fundamental component of visual processing. We frequently use covert attention (i.e., attention that shifts in the absence of eye movements) to preferentially process relevant locations or features in the environment. Measuring the locus and spread of covert attention therefore reveals insights on a wide range of phenomena in visual cognition, such as visual search (Wolfe, 2020), statistical learning (Theeuwes et al., 2022), visual working memory (Oberauer, 2019), reading (Schotter et al., 2012), perception (Carrasco, 2018), and more.

Covert attention is, however, an elusive neural process with no overt marker. Attention research in recent years has therefore focused on developing techniques that track covert attention. Several methods have been added onto the original posner-cueing style behavioural approaches (Posner, 1980) or target-induced ERP approaches (Luck et al., 2000) that have long been used to study attention.

These include microsaccadic gaze shifts (Engbert & Kliegl, 2003; Van Ede et al., 2019), pupillometry (Strauch et al., 2022), and on the neuroimaging side, decoding techniques, such as Inverted Encoding Modelling (IEM) approaches (Samaha et al., 2016). However, these techniques are not without limitations; ideally, to see how attention is distributed across the whole visual field, we would require a tool that provides high temporal resolution, high spatial precision, and the ability to measure attention at several locations concurrently. The above-mentioned tools do not generally meet all these criteria simultaneously.

Frequency tagging approaches offer solutions to these challenges (Norcia et al., 2015). Flickering specific stimuli on the screen at unique frequencies evokes a rhythmic neural response. Modulations to this flicker-evoked response then provide a time-varying measure of (covert) attentional allocation, by quantifying how strongly the corresponding stimuli are being processed. Since frequency tagging is commonly paired with M/EEG, it offers high tem-poral resolution, and since the exact position of the tags is experimenter-defined, it is also spatially very precise. However, presenting multiple low-frequency tags on the screen with unique flickers produces a distracting visual landscape that makes it difficult to distinguish effects of endogenous attention from those caused by the visible flickering of stimuli. More recently, Rapid Invisible Frequency Tagging (RIFT; Arora et al., 2026; Zhigalov et al., 2019) paired with M/EEG has been gaining popularity as an attentional tracker; it extends the classical SSVEP approach, using flickers that are too fast to be perceived. This allows for tracking attentional allocation to multiple regions on the screen, even in the absence of any visible probes that could confound attentional processes.

Theoretically, RIFT could allow us to change our general approach towards localizing spatial attention. Traditional measures of spatial attention such as accuracy derived from behavioural reports, or analyses involving ERPs or endogenous oscillations, often only tell whether attention is or is not directed towards a given location or particular hemifield. RIFT could allow us to instead image attention-driven modulations simultaneously across the entire visual field, rather than probing only a specific location at a time. However, practical constraints currently limit this possibility. RIFT operates within a narrow frequency range (one that permits invisibility while still evoking a neural response). Technical challenges with display equipment often further limit the number of unique frequency tags that may be simultaneously used (Arora et al., 2026). Because of this, current RIFT approaches allow only for sparse sampling of the visual field. To eventually display a large enough number of unique tags to truly map attention across the visual field, we can explore another dimension of tagging selectivity: phase. It has been shown that RIFT responses from simultaneous tags at the same frequency can be isolated in the neural response (Spaak et al., 2024) if they differ in phase. However, it has not yet been determined whether this phase-tagging approach is sensitive enough to be modulated by attention.

Here, in a first, we conduct a RIFT-EEG task with 8 unique tags. We show that RIFT tags can be used to measure attentional-modulations even when differing only in phase, not frequency. This allowed us to develop a basis for the use of combined phase- and frequency-modulated RIFT to paint a picture of attentional allocation across the visual landscape.

## Methods

### Participants

24 healthy participants (age = 22.29 *±* 2.14 years; mean *±* std; sex assigned at birth: 20 female, 4 male) were recruited for this study. 1 participant was excluded due to not being able to fixate properly. The final sample thus consisted of 23 participants. They reported no history of epilepsy and had normal or corrected-to-normal vision. Participants were recruited through Utrecht University’s internal participant framework and were awarded either participation credits or monetary compensation (24€) for participation in the study. The experimental procedure was approved by Utrecht University’s ethical review board of the Faculty of Social Sciences (24-0094). Data were collected between December 2025 and January 2026.

### Protocol

Participants underwent a 2-hour experimental session, prior to which they received information about the experiment and provided informed consent as well as information about their assigned sex at birth, date of birth, and dominant hand. The task procedure was then explained, and the eye-tracker was calibrated. After this participants completed 30 practice trials to get familiar with the task and to practice maintaining fixation. After successful practice run completion, EEG setup followed. The eyetracker was then re-calibrated (and this was repeated at the end of every third block). Participants completed a 60-minute task. The session ended after the successful completion of the task and compensation was awarded.

### Experimental Design & Procedure

Participants performed an orientation discrimination task following an endogenous spatial cue (Figure 1A). The background shape consisted of a circular region divided into eight spatially distinct patches. Each patch displayed a set of lines (referred to as line fields) of which the average orientation could be either +45° or −45° degrees (randomly selected for each patch on each trial). After cue onset, participants reported the orientation of the line field within the cued region, and we used Rapid Invisible Frequency Tagging (RIFT) to uniquely tag each patch and track the allocation of spatial attention over time.

**Figure 1:**
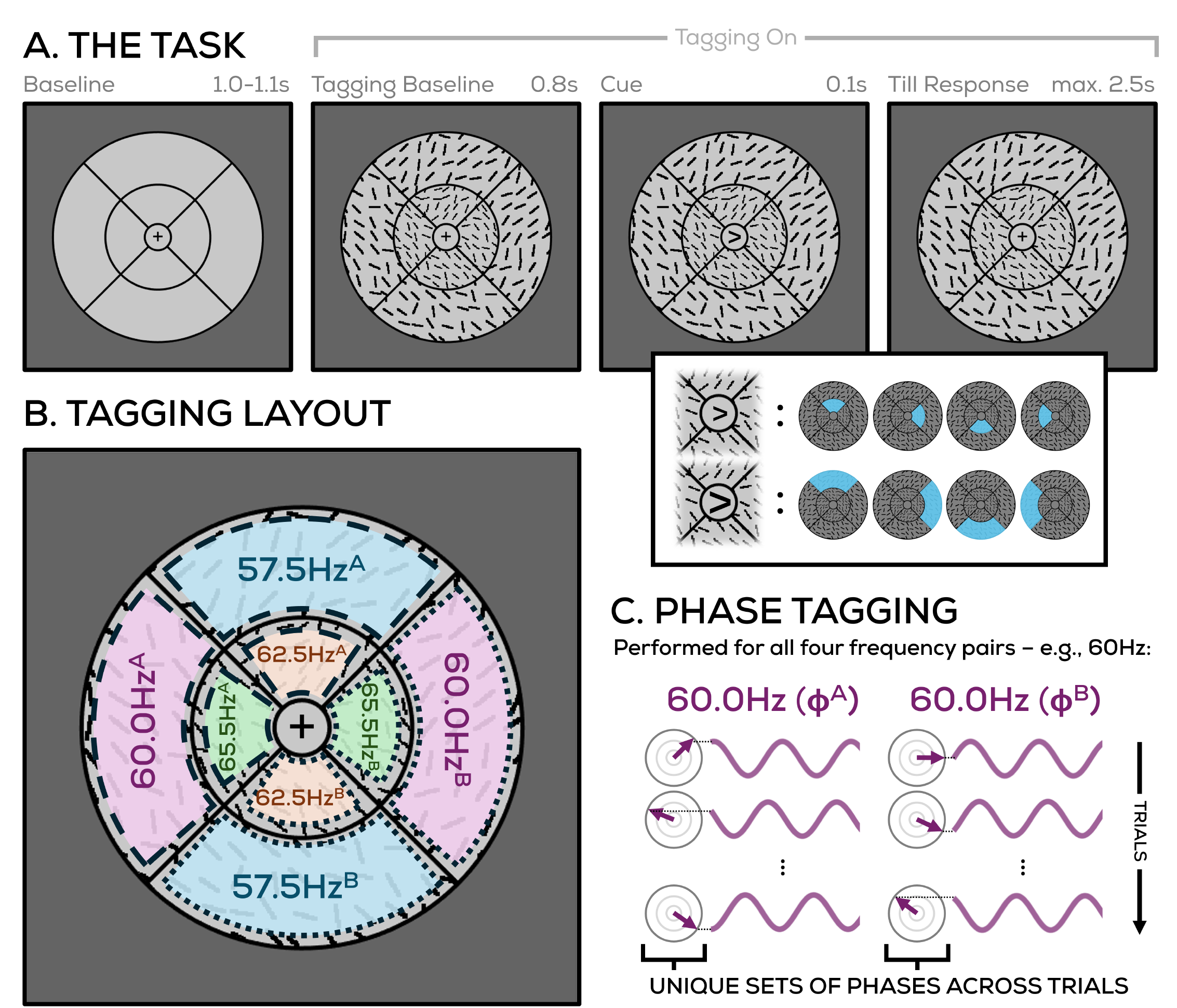
Task Design and Frequency Tagging. A. Participants performed an orientation detection task following a centrally presented spatial cue. Following an initial baseline period, line fields were presented in eight patches across the main stimulus. A centrally presented arrow (based on its size and direction) indicated a cued patch of which the mean orientation had to be reported. Fixation was maintained centrally throughout. **B. Tagging layout**. The luminance of the eight patches was uniquely flickered at four frequencies (65.5Hz, 62.5Hz, 60.0Hz, 57.5Hz), with pairs of tags at the same frequency differentiated by their phase. **C. Phase Tagging**. The two tags at a given frequency were presented with random initial phases on each trial, which gave each tag a unique combination of phases across trials that could later be used to disentangle the two (see *‘Coherence’*).

The task consisted of 15 blocks of 64 trials each. Participants were instructed to maintain fixation on a central cross throughout the experiment, and eye-tracking data was recorded to ensure this. Each trial began with an inter-trial interval (uniformly jittered between 1.0-1.1s) during which tagging was not implemented, line fields were not shown, and an empty silhouette of the stimulus was displayed. Subsequently, line fields were presented in each region of the circular grid. At the same point in time, tagging began as well. This period (lasting 0.8s) is referred to as the “tagging baseline”, since all stimuli (and tagging) had been presented but no attentional cue had yet been shown. This was followed by a central arrow cue (0.1s). It indicated the region to which participants should covertly shift their attention to: its orientation indicated the direction (up, down, left, right) and its size indicated the eccentricity (smaller arrow = inner location, larger arrow = outer location). Participants were instructed to report the average orientation of the line field within the cued region (45° or −45°) by pressing the left or right arrow key. Responses could be made during a 2.5s window following cue offset. Each trial concluded with feedback in the form of a centrally presented check mark or cross for correct or incorrect trials respectively (0.25s). If no response was given or a different key was pressed, the trial was deemed incorrect (median proportion of trials = 0.15%).

### Stimuli

The square background surrounding the main circular stimulus was presented in dark grey (RGB: [80, 80, 80]; see *‘Apparatus’* section for luminance measurements). The main stimulus display consisted of a circular region with an outer diameter of 12 dva, divided into eight regions by a central circular boundary and four radial dividing lines.

The inner circle had a diameter of 6 dva. These regions appeared grey perceptually, but their luminance was varied over time for RIFT (see *‘Tagging Manipulation’*). A circular fixation area of 1.5 dva was presented in grey at the centre of the grid (RGB: [127.5, 127.5, 127.5]), containing a black fixation cross (0.2 dva). All above regions had a black outline (width = 0.08 dva).

Target stimuli consisted of fields of oriented black line elements presented within each of the eight regions of the circular grid. Line fields were generated separately for each of the eight regions. The mean orientation of a given line field was sampled randomly from two possible values (−45° and 45°), and orientations of individual lines were then sampled from a gaussian distribution with this mean value. The standard deviation of this gaussian distribution was varied throughout the experiment as per a staircase procedure (PsychtoolBox QUEST algorithm; Farell and Pelli, 1999; *β* = 3.5, ∆ = 0.01, *γ* = 0.5) running throughout the experiment separately for inner and outer regions. It aimed to maintain task performance around 75% accuracy.

### Tagging Manipulation

Here, we measured the allocation of spatial attention across the screen using Rapid Invisible Frequency Tagging (RIFT). This technique involves flickering the luminance of specific (patches of) pixels faster than can be perceived, and recording the oscillatory response to these luminance changes as a measure of visual processing (or attention).

In this task, we applied RIFT to all eight patches of the circular stimulus region by uniquely modulating their luminance (sinusoidally) over time. We used four tagging frequencies, each obtained by dividing the refresh rate of the PROpixx projector used here (1440Hz; see *‘Apparatus’*) with an integer: 1440/22 (65.45..)Hz, 1440/23 (62.60..)Hz, 1440/24 (60.00)Hz, and 1440/25 (57.60)Hz. This particular set of precise tagging frequencies (hereafter shortened to 65.5Hz, 62.5Hz, 60.0Hz, and 57.5Hz for convenience) was used because they amount to integer divisors of the 1440Hz refresh rate, producing a more consistent tagging envelope. The eight regions of the circular grid were each assigned one of four tagging frequencies, such that each diametrically opposite pair of regions was tagged at the same frequency (Figure 1B). This location-frequency mapping remained consistent throughout the experiment. Thus, outer regions were tagged at 60Hz (left/right) and 57.5Hz (up/down), and inner regions were tagged at 65.5Hz (left/right) and 62.5Hz (up/down).

### Phase Tagging

Conventionally, RIFT studies implement different tags by assigning unique frequencies to each stimulus, so that neural responses can be separated in the frequency domain. To tag a location at a given frequency, we modulated its color from fully black (RGB: [0, 0, 0]) to fully white (RGB: [255, 255, 255]) using a sinusoidal amplitude mask (this mask contained 22, 23, 24, or 25 points per sinusoid cycle for 65.5Hz, 62.5Hz, 60.0Hz, or 57.5Hz respectively). However, here, in addition to tagging different frequencies, we wanted to separate multiple tags at the same frequency. To do so, we also presented each tag on each trial with a random initial phase (selected randomly from the 22, 23, 24, or 25 points on the luminance cycle for the corresponding tag; Figure 1C). These phase offsets were later used during the EEG analysis to differentiate between both tags of the same frequency. Temporal precision of the displayed stimuli was ensured using PsychToolBox’s Screen(‘Flip’) command.

Lastly, to avoid any visible artifacts resulting from onset or offset of the tagging, we smoothly ramped up and down the tagging mask (Minarik et al., 2023). This involved linearly modulating tagging amplitude from 0 to 1 over the first 50ms of tagging onset, and from 1 back to 0 over the last 100ms of tagging onset.

### Apparatus

A ProPixx projector (VPixx Technologies Inc., QC Canada; resolution = 960×540px; refresh rate = 1440Hz) in a rear projection format (48×27.2cm) was used for the experimental display. Luminance was measured to be 0.95cd/m2 for black (RGB: [0, 0, 0]), 649cd/m2 for gray (RGB [127.5, 127.5, 127.5]), and 1330cd/m2 for white (RGB [255, 255, 255]) with an external photometer (Gossen Mavo-Monitor USB) in a dark room. The experiment was programmed and controlled using MATLAB and PsychToolbox (Kleiner et al., 2007). No additional gamma correction was carried out since the ProPixx is already linearized (gamma = 1). To minimize head movements and motor artifacts, participants were stabilized using a chinrest 76 cm away from the screen.

### Eye-Tracking Recording and Analysis

An Eyelink 1000 desktop mount (SR Research, Ontario, Canada) was used to monitor gaze position from both eyes at 500 Hz. At the beginning of the task, a 9-point calibration procedure was performed and was repeated after every third block in the task. Gaze position data was examined to confirm that fixation was maintained centrally during the task. The recorded data were segmented into trials of 1.3s before to 1.5s after cue onset. Trials were baseline corrected with respect to the average gaze position in a 750ms window prior to cue onset. Trials where fixation was not maintained (defined as gaze being further than 1.5 dva from fixation for more than 50ms) were marked. As noted in the *‘Participants’* section, one participant of the 24 collected was excluded from all further analysis due to saccades being made in more than half of the trials. Across the remaining participants, failure to fixate was only observed in a small set of trials (mean = 4.52%, median = 2.40% of trials), and these trials were removed from all EEG analyses.

### EEG recording and preprocessing

EEG was recorded at 2048Hz using 64 active Ag/AgCl electrodes mounted on a fabric/elastic cap from ActiveTwo Biosemi system (BioSemi B.V., Amsterdam, The Netherlands). Two reference electrodes were also placed at the mastoids (but not used). Additionally, the electrooculogram (EOG) was recorded by placing sensors vertically above the left eye and on the left temple. Prior to the start of the task, adequate signal quality was ensured using Biosemi ActiView software. All preprocessing and data analysis was conducted in Matlab using the Fieldtrip toolbox (Oostenveld et al., 2011). The EEG data was first rereferenced to the average of all channels after excluding poor channels determined by visual inspection (median = 12, mean = [mostly frontal] 11.8 channels). Data was high pass filtered (0.01 Hz), then line noise and harmonics were removed using a DFT filter (50, 100, 150 Hz). Next, data was segmented into trials from 1.3s before to 1.55s after cue onset. An ICA was performed to remove oculomotor artifacts, and trials with other (eg. motor) artifacts were manually removed after visual inspection (median = 7.7%, mean = 9.2% of trials) from further analysis. Baseline correction was performed using a 0.4s window (−1.3 to −0.9 s before cue onset).

## Quantifying the RIFT Response

### Coherence

To assess the strength of the EEG response at RIFT frequencies, magnitude-squared coherence (a dimensionless quantity ranging from 0 and 1) was used. Coherence measures the amplitude of a given frequency in a given input signal, but also measures how strongly this frequency component is phase-locked over trials. This emphasis on phase-locked signals forms the basis for how we separated distinct tags at the same frequency.

Here, we had randomized the phase onsets of each tagging signal across trials (see *‘Tagging Manipulation’* section). This required us to subsequently realign the EEG data to the specific trial-wise phases of any chosen tagged region, because coherence considers phase-locked responses. The EEG response to that tagged region would then be phase-aligned across trials, while responses to other tagged regions, including those sharing the same frequency but carrying a different phase, would not be phase-aligned. Computing coherence on this phase-aligned EEG response then yields a response specific to the tagging region corresponding to that set of phases, without any contribution from other locations. This enabled independent reconstruction of neural responses from all eight tagged locations despite their simultaneous implementation.

We first explain the basic coherence computation, after which we explain how it was implemented for the different analyses used here. Segmented and phase-aligned trials were bandpass filtered at the frequencies of interest using a two pass Butterworth filter (4th order, hamming taper, bandpass width = *±* 1.9Hz). Magnitude-squared coherence is computed using a reference signal, for which we used pure sinusoids at the corresponding frequencies. The filtered EEG response and reference sinusoid were Hilbert-transformed to obtain their analytic signals, from which time-varying instantaneous magnitude (*M* (*t*)) and phases (*ϕ*(*t*)) were extracted. These instantaneous magnitudes and phases were used to compute time-varying coherence as per Equation 1 below, where *x* and *y* subscripts represent EEG signal and reference signal metrics respectively. This procedure was carried out independently for all EEG channels.

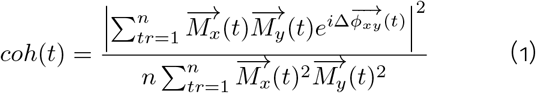

### Applying coherence to the data

Coherence was used for three different analyses, (1) confirming that all 8 tagging responses could be separately recovered (Figure 2), (2) determining whether attentional differences could be measured between tags of the same frequency (Figure 3), and (3) quantifying attentional allocation across the visual field after the cue (Figure 4). For all these analyses, coherence was averaged over the six channels that showed the strongest mean coherence during the tagging interval for each participant and tag. This approach is commonly used (Dietz et al., 2025; Hustá et al., 2025; Wang et al., 2026), and it has been shown that the specific choice of number of channels does not affect the observed attentional modulation (Arora et al., 2026).

**Figure 2:**
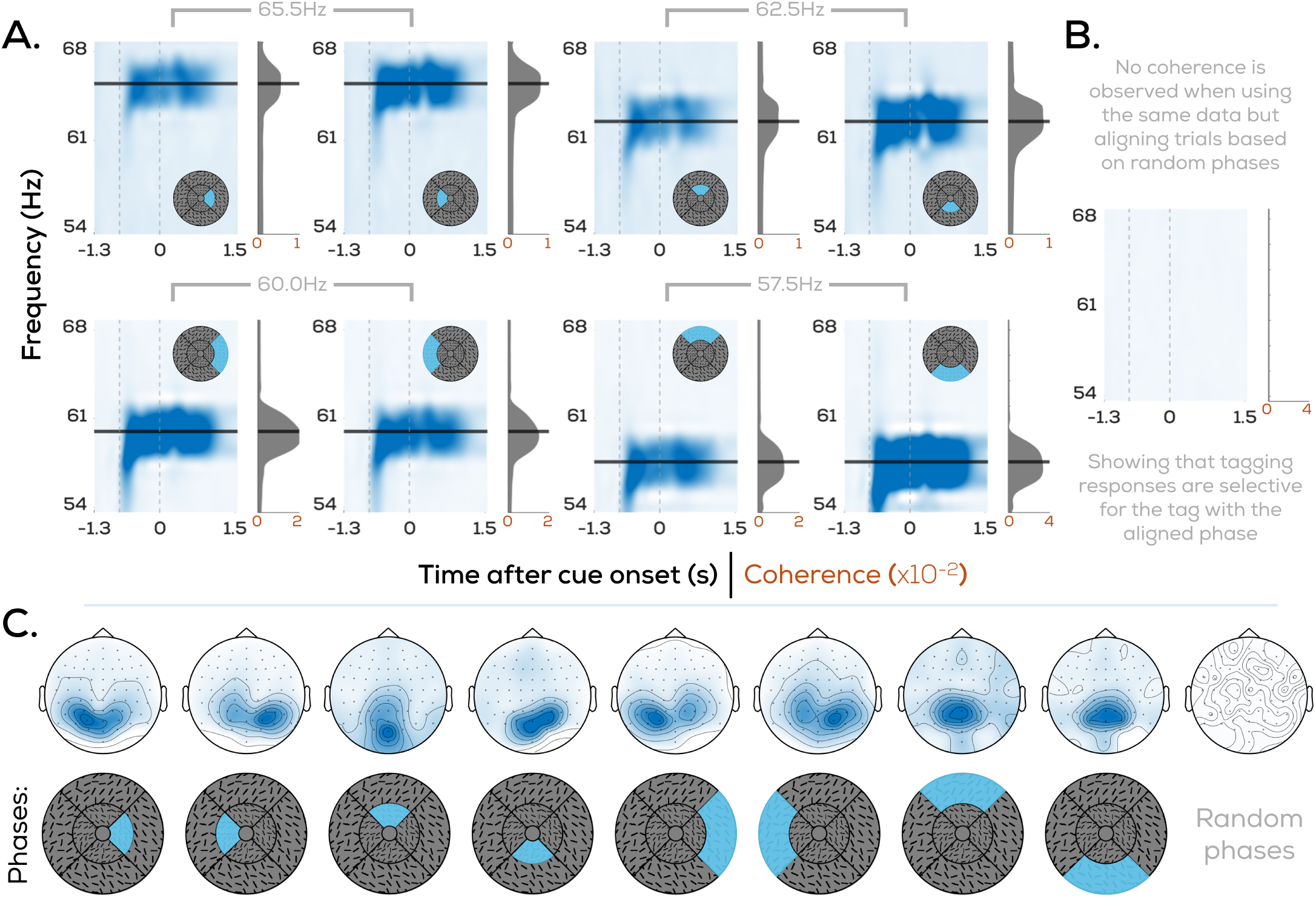
Tagging responses from eight simultaneous RIFT tags. Spectrograms showing mean coherence in the 54-68Hz frequency range over the course of a trial when phase-aligning the EEG data to **A**. phases corresponding to each of the 8 tagging locations, **B**. random phases. Vertical, dotted lines represent tagging onset (t=-0.8s) and cue onset (t=0s) respectively. Horizontal, solid lines mark the stimulated frequency for the corresponding tag. **C**. Topographical spread of mean coherence for each of the 8 tagging locations (averaged from t=-0.75s to t=1s after cue onset). Coherence topoplots are plotted with unique colour axis limits; color axis ranges from 0 (minimum) to [0.03, 0.05, 0.03, 0.05, 0.12, 0.07, 0.06, 0.20, 0.05] (maximum) for each of the 9 topoplots from left to right respectively.

**Figure 3:**
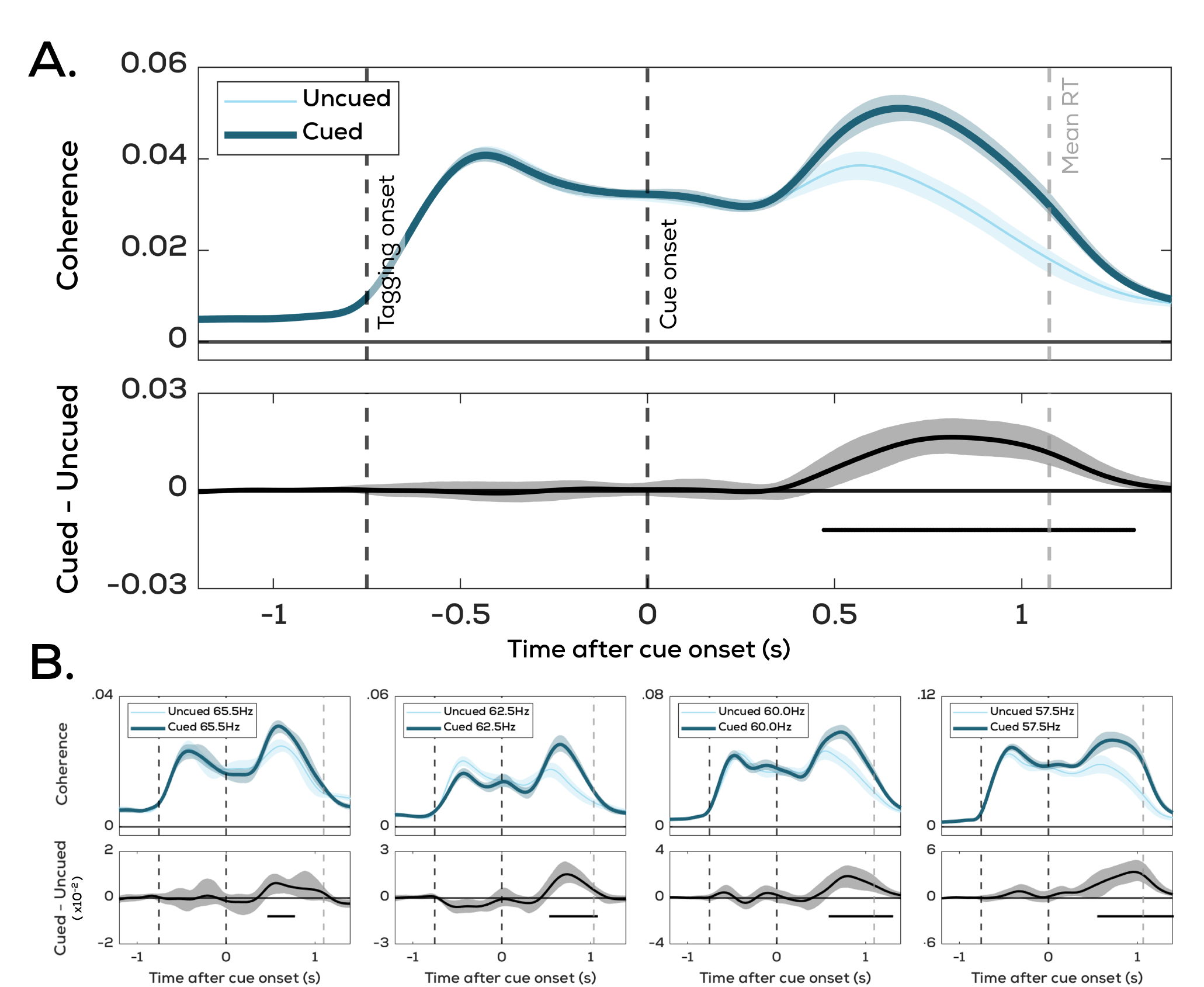
RIFT Responses reflect the cued tagging location. Cued and uncued coherence traces over the trial compared **A**. for the average, and **B**. individually for the four pairs that were tagged at the same frequencies. Shaded regions represent 95% bootstrapped confidence intervals (cousineau-corrected; Cousineau, 2005). Horizontal bars indicate significance as per permutation cluster testing. Horizontal dotted lines reflect tagging onset, cue onset, and mean reaction time respectively.

**Figure 4:**
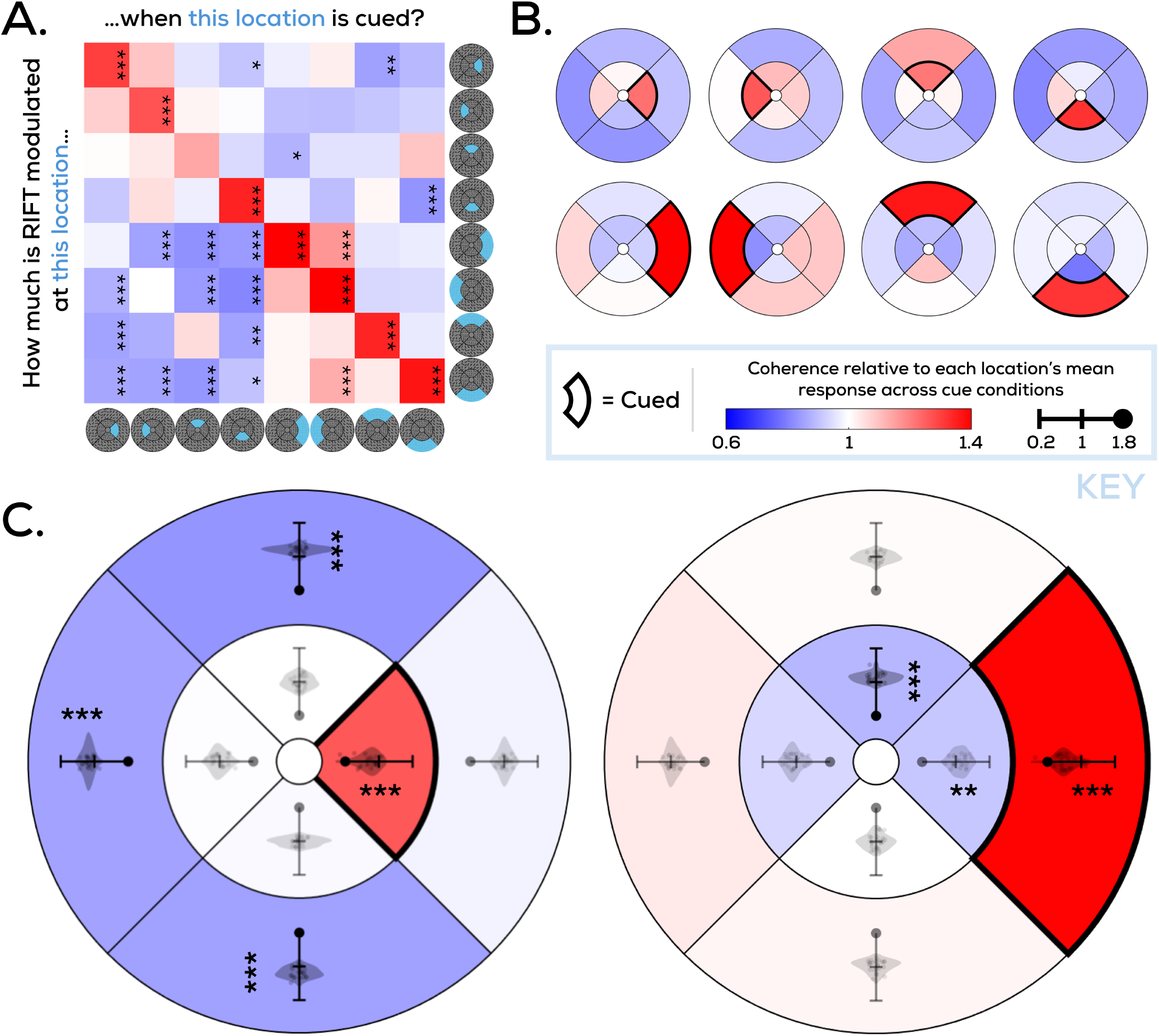
Attentional allocation across the visual field. Mean coherence modulations across participants as computed for all eight tagging regions separately over sets of trials where each region was cued, **A**. visualized in tabular format, and **B**. overlaid onto the main stimulus shape. For all panels, the color axis in ‘Key’ applies; values of 1 represent no change compared to mean responses of the corresponding location across cue conditions, values above/below 1 represent increased/decreased RIFT responses. **C**. Data in A-B averaged across inner cue conditions and outer cue conditions (by rotating the plots in B. to produce cue-locked figures). Axes within the tagged regions reflect distributions of the corresponding values across participants; significance determined as per 95% bootstrapped CIs. Stars represent significance (1: 0.025 < p < 0.01; 2: 0.01 < p 0.001; 3: p < 0.001).

For constructing spectrograms and topographical distributions of the tagging response, coherence was computed for frequencies between 53.6Hz and 68.8Hz in 0.8Hz intervals over the whole trial window. Topographical distributions were produced by averaging the interval from −0.75s to 1s after cue onset. In this case, data was phase-aligned (see *‘Coherence’*) to each tag one-by-one. For example, phase aligning the data using the trial-wise phases of the 60Hz tag on the left side of the stimulus display will produce the tagging response specifically (and only) for this left-side 60Hz tag.

Next, we looked at attention-induced differences in the tagging response, by comparing RIFT coherence between cued and uncued tags of the same frequency. Since we only looked at differences between the pairs of tags that flickered at the same frequencies (but differed in phase), we first split the trials based on whether a region tagged at 65.5Hz, 62.5Hz, 60.0Hz, or 57.5Hz was cued. As explained above, phase-aligning the EEG data to the phases of a particular tagged region allowed us to isolate the tagging response of that specific region without any contribution from other tagged regions. Alternatively, we can also align the data to the trialwise phases of the “cued” 60Hz tag, regardless of whether the cued tag happened to be the one on the left or the right. Similarly, we can align the data to the trialwise phases of the “uncued” 60Hz tag. This way we get two coherence traces at the same frequency, from the same trials, differing only in whether we obtain this signal from a tag that had been cued or not. We repeated this process for all four tagging pairs at matching frequencies, producing four cued vs. uncued coherence traces per participant. These were later averaged together to produce one cued vs. uncued set of coherence traces per participant.

### Exploratory Analyses to quantify the Distribution of Attention around the Visual Field

The main goal of our experiment was to test whether we could measure attentional effects at regions that were tagged at the same frequency but varied in phase on a trial-by-trial basis. In addition to this, we also performed an exploratory analysis of how attentional allocation varied across the whole display when a particular region was cued, since our eight tags spread across the visual field allowed us to do so. However, we had not designed our experiment to test specific hypotheses about the precise distribution of attention. Thus, we limited ourselves to visualizing our results and statistically estimating if a tagged region showed higher or lower RIFT responses compared to its average across cue conditions. Notably this means that our assessment of attention following the cue (as indexed by coherence) is relative and should not be interpreted as suppression. More specifically, this follows from a limitation inherent to our analysis: we cannot directly use the coherence traces obtained from each location, since different tagging frequencies and tagging locations have different overall tagging signal levels. This required us to perform a normalization of the data to make tagging signals from various locations comparable. The precise choice of the normalization can affect the resulting attentional distribution and how it is interpreted. As a thorough exploration of the effects of these analyses choices was beyond the scope of this paper, we limited ourselves to an exploratory visualization and basic statistical assessment which are included in the discussion.

As described above, we quantified how attention was distributed across the whole visual field (i.e., all eight regions) when a particular region was attended. To do so, we split the trials into eight sets depending on which location was cued, and ensured at this stage that all eight cue conditions had an equal number of trials per participant by removing random trials from conditions that had retained more trials after preprocessing. With 8 tagging locations, and 8 possible cue conditions, we produced an 8×8 set of coherence traces (phase-aligning to each of the 8 locations when each of the 8 locations was cued). We then performed two preliminary steps before presenting this data. First, to obtain a time-averaged version of the cue-induced attentional modulation, we averaged coherence over the interval during which a significant attentional effect was seen at the group level in the previous analysis (0.5s-1.2s). Secondly, to avoid any effects of overall fluctuation in coherence between various tagging locations (e.g. due to the known effects of tagging frequency, eccentricity, and visual field asymmetries on the tagging response; Arora et al., 2026), we divided the above-produced time-averaged values within each tagging location, with their means across the various cue conditions. That is, if a particular tagging location’s coherence response is relatively lower regardless of the cue condition, these effects are averaged out, and we only look at the relative difference between cue conditions.

In this manner, we obtained one value reflecting cue-induced coherence modulation for each tagging location per cue condition. We then separately averaged conditions where an inner location was cued, or an outer location was cued (while rotating them by the corresponding angle so that one “cue-locked” result was produced). When producing a time-varying version of the same figure (Supplementary Video 1-2), all steps followed were the same as those above, with the exception of applying baseline correction to each timepoint rather than to the average coherence signal from the interval 0.5s-1.2s.

### General Statistical Analysis

When comparing the tagging response coming from cued vs. uncued tags, we obtained one coherence trace corresponding to cued and uncued tags per participant. This was the case both when looking at individual frequency pairs, or at the overall comparison averaged across frequencies. By subtracting the uncued trace from the cued trace, we obtained one difference trace per participant. These difference traces were tested for significance while accounting for multiple comparisons using a cluster permutation test (Maris & Oostenveld, 2007), the steps of which were as follows: first, a one-sample t-test was run for each individual timepoint of the set of difference traces to detect points on the trace significantly different from 0 (p < 0.1). Any clusters of consecutively significant time-points (minimum 10ms long) were selected, and a cluster t-mass was obtained for each of these by summing the t-values of each point in that cluster. This cluster t-mass was tested for significance by comparing it to a bootstrapped distribution of cluster t-masses. The bootstrapped distribution was obtained by computing 5000 iterations where we randomly flipped the sign of each individual difference trace (to preserve the temporal correlation of the traces), and added the highest cluster t-mass to the distribution. Lastly, a cluster from the original difference traces was labelled as significant when its t-mass was in the bottom or top 2.5% of cluster t-masses within the bootstrapped distribution.

To determine whether the time-averaged coherence modulations in Figure 4 were significant, we computed bootstr--apped distributions for the 23 (one per participant) values per tagging location per cue condition. Significant modulation was inferred if the bottom or top 2.5% of the boot-strapped distribution included zero.

## Results

Participants successfully performed the task (mean accuracy = 80.8%, 95% bootstrapped CI = [79.3%, 81.6%]) well above chance level (50%). This was similar for trials where the inner (mean = 79.8%, 95% bootstrapped CI = [78.0%, 81.0%]) and outer (mean = 81.7%, 95% bootstrapped CI = [80.5%, 82.4%]) locations were cued (two staircases were run in parallel for the outer and inner line field stimuli, see *‘Stimuli’*).

We first examined whether we could successfully evoke measurable RIFT responses for each of the eight tagged regions. On all trials of the experiment, all eight tags were simultaneously presented. Coherence analysis nonetheless revealed clear frequency-specific peaks for each tagged region at its corresponding tagging frequency (Figure 2A). We isolated responses from these tags only by changing which set of phases (corresponding to 1 of the 8 tags at a time) the trialwise phase-alignment was based upon (see *‘Quantifying the RIFT Response’*). Had this phase-alignment method been insufficient in its tag-selectivity (i.e., had it resulted in leakage of RIFT response across tags), this would have resulted in additional oscillatory peaks (e.g. a 60.0Hz peak when phase-aligning to 57.5Hz tags). The absence of such additional peaks confirms our ability to selectively measure responses from only one of our tags at a time, even those that share a tagging freque--ncy. We can illustrate this better by using a random set of phases across trials that does not correspond to any of the eight tags (Figure 2B); in this case, no coherence is observed at any of the tagged frequencies. Consistent with the previous RIFT-EEG literature (Arora et al., 2025; Bouwkamp et al., 2025; Dimigen et al., 2025; Spaak et al., 2024; Zhigalov et al., 2019), coherence was topographically the highest over parietal and occipital electrodes across all locations (Figure 2C).

Next, we looked at cue-induced attentional modulations to the RIFT response. Specifically, we compared RIFT responses between tags of each frequency pair when one of them was cued. Coherence traces corresponding to mean cued vs. uncued frequency tags showed a clear difference emerging after cue onset (Figure 3A; significant cluster from ~0.5-1.3s after cue onset; p < 0.0002; cluster t-mass compared with bootstrapped distribution; note that the cluster permutation test used here specifically tests whether an effect is present or not; it is not a measure of effect latency). The 95% CIs of the attentional modulation departed from zero well before the mean RT (~650ms), implying that it is not in some way caused by the manual response. Prior to averaging all frequency pairs together, all four individual frequency comparisons also showed the same cued vs. uncued difference Figure 3B: 65.5Hz (~0.5-0.7s after cue onset; p = 0.0236), 62.5Hz (~0.5-1.1s after cue onset; p = 0.0006), 60.0Hz (~0.6-1.3s after cue onset; p = 0.0016), and 57.5Hz (~0.5-1.5s after cue onset; p = 0.0002). This shows that using phase, attention-induced differences in tagging responses can be measured even between two stimuli tagged at the exact same frequency.

## Discussion

Here, we show that Rapid Invisible Frequency Tagging (RIFT) can be combined with phase-alignment to record neural responses corresponding to at least 8 simultaneously tagged locations. We also show that phase can be used as an additional tagging feature to measure attentional differences even between regions tagged at the same frequency. Overall, we demonstrate the feasibility of phase-aligned RIFT as a method for tracking how spatial attention is distributed across the visual field.

### Phase Tagging can be used to measure covert attentional differences

We observed clear and distinguishable tagging responses from all eight simultaneous locations individually on the screen. This is the highest number of concurrent tags implemented in a RIFT task so far, and a key feature in this implementation was the usage of phase as a tagging feature.

RIFT operates within a narrow frequency band where the tagging is too fast to be perceived, but still slow enough to evoke a response in early visual cortex (Duecker et al., 2025; Ferrante et al., 2023; Minarik et al., 2023). This places an upper limit on the number of frequency tags that can be feasibly implemented without responses from adjacent frequencies overlapping. Given that such a limit exists when using one tag per frequency, using phase as an additional tagging feature (i.e., having more than one tag per frequency) naturally increases this limit. Phase tagging could therefore enhance our ability to track the locus of spatial attention over time, as exemplified here with our visual field maps of attentional allocation. Measuring attentional modulations across the entire visual field simultaneously can allow us to think of attention more broadly than as something that is “present” at one given location or acts on one given stimulus relative to one comparison point, and towards creating a holistic overview of how attention is allocated across the visual field.

We combine RIFT and phase-tagging here by assigning both tags (of the same frequency) a random initial phase on each trial. Theoretically, this procedure is not limited to only two phase tags; any number of tags at the same frequency could be implemented in this manner, as long as there are enough trials to make the random phase combinations sufficiently unique. This is one of the ways in which our design differs from a previous implementation where phase tagging was combined with RIFT during passive viewing (Spaak et al., 2024). Spaak and colleagues implement RIFT-phase-tagging by maintaining a constant phase difference between the tags across trials (90°). Assigning different tags specific frequencies, such as orthogonal, may place a limit on how many such phase tags may be used, since phases infinitesimally close together would be difficult to separate (some SSVEP-based phase tagging setups have suggested eight tags as an upper limit; Tong and Zhu, 2015). Our approach, which instead relies on the unique sequence of phases each tag has across trials, may be more suited to implementing a larger number of tags at the same frequency, without having to consider whether consistent phase-differences are too close together. However, since different tags can have similar phases on a given trial, our approach may make it difficult to identify attended tags at the single-trial level. Future studies should ideally ensure that different tags of the same frequency do not have the same phase on a given trial. Our approach also does not require collecting any preliminary data, allowing for an easier experimental implementation as compared to the existing approach. There, RIFT responses from a stimulus tagged with random phases over trials are first used to estimate a tagging-to-brain phase lag, which can then recover independent RIFT responses from two simultaneous tags at different phases (Spaak et al., 2024).

The idea of introducing phase as a feature for frequency tagging is not original to RIFT studies, but has been explored earlier with traditional SSVEP studies. These studies were typically designed to measure overt gaze responses in order to develop Brain-Computer Interfaces (Jiang et al., 2019; Mao et al., 2020; Wittevrongel and Van Hulle, 2016; Yeh et al., 2013; but see Armengol-Urpi et al., 2021 for one exception aimed at tracking covert attention). Regardless of the application, similar to Spaak et al., 2024, prior SSVEP studies have implemented phase tags by assigning a constant phase offset to different tags relative to each other, rather than using random phases as we do here. Here, we instead recover tagging responses from tags by simply skewing our EEG data slightly along the time axis to phase-align the responses of a particular tag. This general procedure is far more feasible with RIFT rather than traditional SSVEP, because skewing trials in this manner naturally introduces some temporal smoothing into the data, but the extent of this smoothing depends on the tagging frequency. As opposed to traditional SSVEPs, the high frequencies used for RIFT produce negligible amounts of such smoothing (<8ms, Arora et al., 2026). Thus, RIFT allows us to use phase-alignment of tags with random phases with minimal concerns over temporal smoothing.

This study represents a first step towards using phase in combination with invisible frequency tagging to track the distribution of covert attention across the visual field. Future work may use this implementation as a stepping stone to explore how phase and frequency tagging can be further optimized to increase the number of dissociable tagging responses.

### Exploratory Analyses Quantify the Distribution of Attention around the Visual Field

In the attentional comparisons of Figure 3, we tested our main hypothesis by contrasting the RIFT response between the cued and uncued locations of a particular shared tagging frequency pair that differed in phase. However, the eight tags (varying in frequency and phase) that we implemented here could theoretically also be used to assess how attentional allocation varied across the whole display when a particular region was cued. Methodologically this would allow researchers to test a variety of hypotheses about how attention is distributed dynamically with high spatial resolution. In the following section we present results from an exploratory analysis where we visualized the spatial distribution of attention in all cueing conditions. Our aim was to provide a preliminary and exploratory demonstration of the potential of our methodological contribution rather than test specific hypotheses. Crucially, because RIFT responses differ across tagging frequencies and locations, comparisons across the display require normalization, the choice of which can influence the resulting patterns. The following results should there-fore be interpreted as relative changes in RIFT responses across cue conditions, rather than as direct measures of attentional enhancement or suppression at individual locations.

To do so, we computed one ‘modulation’ value for each tagging location, reflecting how much the RIFT response was modulated after the cue. This was done separately for each of 8 cue conditions (Figure 4A-B). Note that these values are baselined with respect to the mean RIFT response measured at that location, across all 8 cue conditions (see the section *‘Quantifying the RIFT response’* for a detailed description on how these time-averaged values were obtained). Thus, positive values reflect a relatively higher RIFT response at that location following the cue, and negative values reflect relatively lower responses. The resulting maps were then rotated, so that the cued location was always on the ‘right’ and averaged across all four conditions where an inner (Figure 4C, left) or outer (Figure 4C, right) region was cued to produce a “cue-locked” map visualizing how the RIFT response changed in response to the cue, simultaneously across all 8 tagged locations.

When outer tagging regions were cued, as expected, we observed a positive coherence change at the cued location (mean modulation: +0.398, p < 0.0002), but we also observed a negative coherence change at the inner location directly adjacent to this cued location (−0.085, p = 0.0056), and the inner location counter-clockwise from the cued outer region (−0.110, p < 0.0002). When inner tagging regions were cued, we again observed a positive coherence change at the cued location (+0.264, p < 0.0002). Surprisingly, in this case we also observed a negative coherence change at all outer tagging regions that were not adjacent to the cued region (outer region clockwise from the cued inner region: −0.146, p < 0.0002; outer region diametrically opposite to the cued inner region: −0.142, p < 0.0002; outer region counter-clockwise from the cued inner region: −0.163, p < 0.0002). These patterns are also observed in all of the individual cue conditions (Figure 4A-B), showing that the pattern of results obtained with the average maps is not driven by a peculiar response in a single cue condition.

Lastly, to facilitate intuitive interpretation of the spatio-temporal evolution of attention following a cue, we compiled the RIFT responses across all eight locations (relative to the cued location, as depicted in Figure 4) in a comprehensive video (Supplementary Video 1-2). Though we do not reflect on the theoretical mechanisms of such a distribution of attention extensively here, the results seem to suggest that attention may be distributed differently when lower and higher eccentricities are cued. Further testing may elucidate the source of these differences. Importantly, these exploratory analyses demonstrate that RIFT can be used to investigate the spread of attention across the visual field.

## Conclusion

We showed that phase-modulation can be incorporated into RIFT experiments to measure attentional differences in visual processing from at least 8 simultaneously tagged locations. We also demonstrate the feasibility of phase-aligned RIFT as a method for tracking how spatial attention is distributed across the visual field.

## Supplementary Material

Links to Supplementary Videos 1 and 2:

1. Supplementary Video 1 (Inner)
2. Supplementary Video 2 (Outer)

**Supplementary Video 1-2: Attentional allocation across the visual field visualized over time for the inner (1) and outer (2) cue conditions**. Here, we present the same distributions as those of Figure 4C, but over time. Values of 1 represent no change compared to mean responses of the corresponding location across cue conditions, values above/below 1 represent increased/decreased RIFT responses. Timestamp in the top-right corner represents time after cue onset.

## Data and Code Availability

Raw and Preprocessed EEG data is available at: https://osf.io/f47wc/overview

## Declaration of Interests

The authors declare no competing interests.

## Funding Declaration

This project was supported by a VICI Grant awarded to S.V.d.S. by the Netherlands Organization for Scientific Research.

## Author Contributions

Conceptualization, K.A., L.K., S.V.d.S., S.G., S.C.; Data Curation, K.A., K.Z.; Formal Analysis, K.A., K.Z., S.C.; Funding Acquisition, S.V.d.S.; Investigation, K.A., K.Z.; Methodology, K.A., S.G., S.C.; Project Administration, K.A., S.C.; Resources, S.V.d.S.; Software, K.A., S.C.; Supervision, L.K., S.V.d.S., S.G., S.C.; Visualization, K.A.; Writing – Original Draft Preparation, K.A., K.Z., S.G., S.C.; Writing – Review & Editing, K.A., K.Z., L.K., S.V.d.S., S.G., S.C.

## Notes

### Competing Interest Statement

The authors have declared no competing interest.

